# Efficient production of itaconic acid from the single carbon substrate methanol with engineered *Komagataella phaffii*

**DOI:** 10.1101/2024.04.25.591069

**Authors:** Manja Mølgaard Severinsen, Simone Bachleitner, Viola Modenese, Özge Ata, Diethard Mattanovich

## Abstract

**Background:** Amidst the escalating carbon dioxide levels resulting from fossil fuel consumption, there is a pressing need for sustainable, bio-based alternatives to underpin future global economies. Single carbon feedstocks, derived from CO_2_, represent promising substrates for biotechnological applications. Especially methanol is gaining prominence for bio-production of commodity chemicals.

**Results:** In this study, we show the potential of *Komagataella phaffii* as a production platform for itaconic acid using methanol as the carbon source. Successful integration of heterologous genes from *Aspergillus terreus* (*cadA*, *mttA* and *mfsA*) alongside fine-tuning of the *mfsA* gene expression, led to promising initial itaconic acid titers of 28 g·L^-1^ after five days of fed-batch cultivation. Through the combined efforts of process optimization and strain engineering strategies we further boosted the itaconic acid production reaching titers of 55 g·L^-1^ after less than five days of methanol feed, whilst increasing the product yield on methanol from 0.06 g·g^-1^ to 0.24 g·g^-1^.

**Conclusion:** Our results highlight the potential of *K. phaffii* as a methanol-based platform organism for sustainable biochemical production.

## Introduction

Due to the continuous exploitation of fossil fuels, the atmospheric content of carbon dioxide is rapidly increasing [1], and it becomes clear that alternative, sustainable, bio-based products and feedstocks must form the foundation of the global economy in the future. Efforts are underway to pioneer novel technologies and production platforms for products traditionally produced from gas, coal or oil [2,3]. Many of these advancements use microbial systems to enable conversion of organic materials into biofuels as well as chemicals with a wide range of complexity [4–8]. The replacement of fossil resources with renewable first-generation feedstocks, however, raises social and ethical concerns due to the high demand for alternative food and feed sources driven by the growing world population [9,10]. Naturally, this has prompted greater technological investment in alternative substrates, aiming for a neutral carbon footprint and sidestepping ethical implications. Sustainable resources of interest include various waste products such as crude glycerol [11], lignocellulosic biomass [12], food waste [13] and more unconventionally single carbon (C1) substrates derived from CO_2_ [14].

Valorising CO_2_ and its derived C1-substrates is a growing area of interest, being developed in various biotechnological [4,15–17] and chemical fields [18]. Methanol (MeOH), a low-cost renewable single-carbon feedstock, is gaining attention for bio-production of commodity chemicals. With the additional interest in MeOH, production capacity increased to 174 million metric tons by 2022 and is steadily increasing [19]. Traditionally sourced from syngas, MeOH can now also be generated from methane and CO_2_, offering a means of sequestering greenhouse gasses to support a sustainable bio-economy [20]. Methylotrophic organisms, possessing the inherent capability to utilize MeOH as their exclusive carbon and energy source, stand out as promising candidates for platform organisms, with the yeast *Komagataella phaffii* being one of them [20–22]. Although *K. phaffii* is widely recognized in the industry for recombinant protein production [23], recent findings indicate its significant potential in generating platform chemicals [24–27].

Itaconic acid (IA), an unsaturated dicarboxylic acid, holds considerable promise as a biochemical building block, as its versatility lies in its ability to serve as a monomer for various products such as resins, plastics, paints, and synthetic fibres [28,29]. Traditionally, microbial production of IA started in 1960 by fermenting *A. terreus* in sugar-containing media [30]. Other microorganisms like *Ustilago* sp., *Candida* sp. and *Rhodotorula* sp. have since shown the ability to produce IA, however, *A. terreus* has remained the predominant industrial production host, achieving titers of up to 150 g·L^-1^ during a cultivation for 9.7 days [13,31–33]. The natural biosynthetic pathway for IA follows a pathway analogous to citric acid formation, involving the tricarboxylic acid cycle (TCA). In *A. terreus*, IA is produced from cis-aconitate by the key enzyme aconitate decarboxylase, CadA, residing in the cytosol [34–36]. Further two transporters are involved in shuttling either the precursor cis-aconitate from the mitochondria to the cytosol, i.e. the mitochondrial cis-aconitate transporter (MttA) or the product itself to the extracellular space, i.e. the Major Facilitator Superfamily transporter (MfsA) [36,37].

In this study we explored the potential of *K. phaffii* as a production organism for IA using MeOH as the primary carbon source. For that, the heterologous genes *cadA*, *mttA* as well as *mfsA* from *A. terreus* were successfully engineered into *K. phaffii.* Upon small-scale preliminary experiments in 24-deep well plates and shake flasks, fed-batch cultivations were carried out in lab-scale reactors, where optimal results were achieved by combining strain engineering with process engineering.

## Methods

### Cloning and yeast transformation

The *Komagataella phaffii* wild type strain CBS7435 (Centraalbureau voor Schimmelcultures, NL) was used as a host for engineering the initial itaconic acid (IA) producing strain, the cadA strain. In CBS7435 a single copy of the cis-aconitate decarboxylase gene from *A. terreus*, *cadA* (ATEG_09971) was genomically integrated under the control of the p*AOX1* in the *RGI2* locus using CRISPR-Cas9. In a similar manner, the cadA+mttA strain was generated, where both transcription units, *cadA* and *mttA* (ATEG_09970), were incorporated into the RGI locus in CBS7435. For *mttA* expression the p*POR1* promoter was used. For the cadA+mttA+mfsA strains, homologous recombination of the *mfsA* gene (ATEG_09972) encoding a cytoplasmic IA exporter was inserted into the *GUT1* locus by CRISPR-Cas9 in the cadA+mttA strain. To find the optimal expression strength of *mfsA*, different promoters (pGAP, p*FDH1*, p*POR1*, p*FBA1*) were tested. The cadA+mttA+mfsA_pGAP_ strain was used as host for generation of the multicopy strains (MC). To allow random integration of heterologous genes via non-homologous end joining, plasmids of the respective genes were created with unspecific homologous overhangs and without the assistance of CRISPR-Cas9. dDNA was prepared by linearization using AscI and PCR purification (Jena Bioscience PP-201L). Transformations were facilitated in a single round to generate strains with multiple copies of the individual genes; of *cadA* and *mfsA;* and of all three genes simultaneously. The transformed multicopy clones were plated and selected on plates with high zeocin concentrations (300, 600, 1000 µg·mL^-1^). Integrations of donor DNA (dDNA) into correct loci or random integration were verified via PCR for all clones. A list of the strains used in this study are listed in Table 1.

**Table 1:**
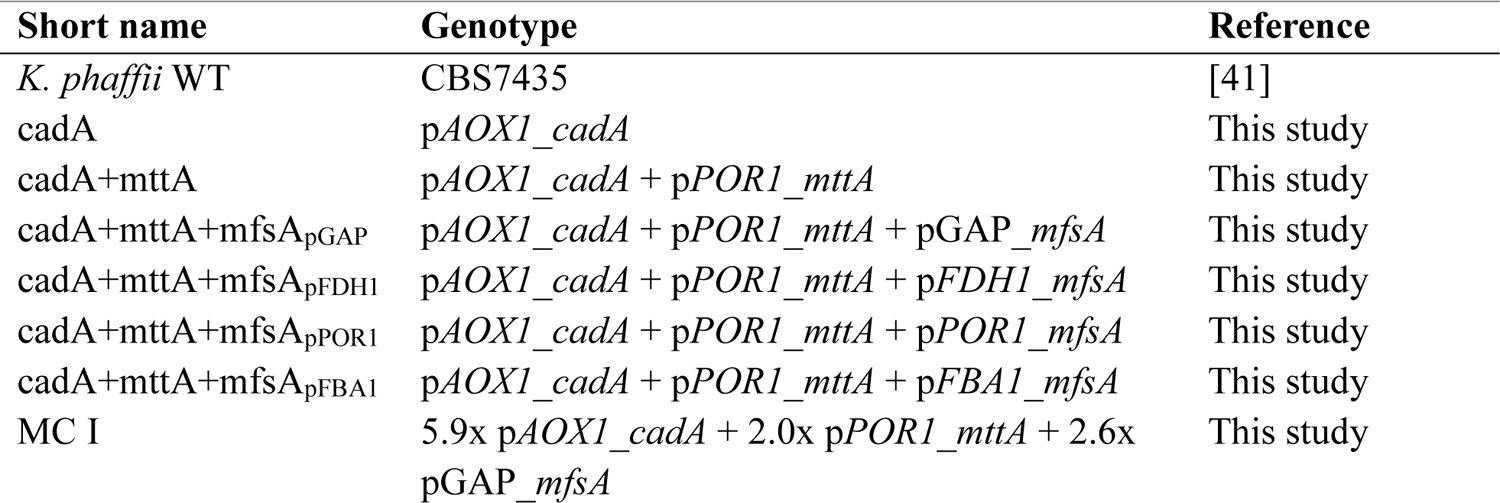
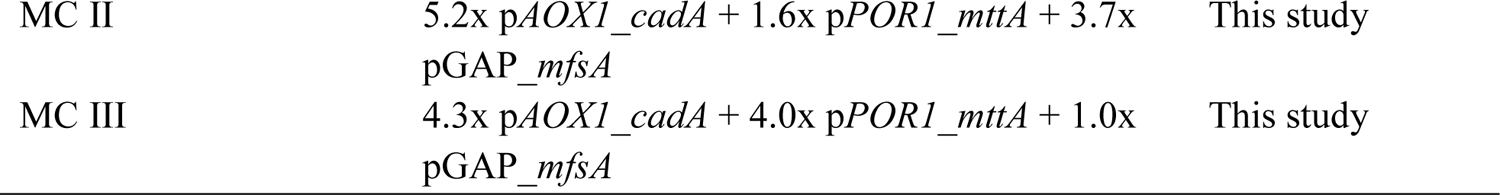
*K. phaffii* strains used in this study.

The dDNA, consisting of the respective heterologous genes flanked by the homologous sequences of the respective integration loci, was prepared by amplification by PCR using Q5® High-Fidelity DNA Polymerase from the respective plasmids. Primers and plasmids were designed using Qiagen CLC Genomics Workbench. Primers were ordered at Eurofins Genomics, sequences and names can be found in Table S1. All plasmids were designed using the Golden Gate Assembly (GGA) and the CRISPR-Cas9 systems developed and described by [38,39], utilizing *Escherichia coli* DH10B as host. *E. coli* were transformed with the respective plasmids via heat-shock, transformants were then grown on LB plates with either 50 µg·mL^-1^ kanamycin or 50 mg·mL^-1^ ampicillin [38,39].

Transformation of *K. phaffii* was performed as described by [40]. To summarize, 80 µL of competent *K. phaffii* cells were gently mixed with the linearized DNA and the CRISPR-Cas9 vector [38,39], the mixture was then transferred to a chilled electroporation cuvette for 5 min, and electroporated with BioRad Gene Pulser^TM^ for 4 ms (2000 V, 25 µF, 200 Ω). The transformed cells were then mixed with cold YPD and regenerated for 2 hours at 28°C before plating on YPD or YPG plates with respective antibiotics (kanamycin or zeocin) and incubation at 30°C for two days.

### 24-deep well plate screenings

First screenings to identify producing strains were performed in 24-deep well plates of 2 mL cultures for 48 h. Here a two-phase screening protocol was used, as described by [40]. The first phase consisted of biomass production, where selected clones were cultivated in 2 mL YPG precultures overnight at 25°C or 30°C and 280 rpm. The main cultures were initiated by inoculating buffered Yeast Nitrogen Base (YNB; 3.4 g·L^-1^, pH 5.5-6.0, supplemented with 10 g·L^-1^ (NH_4_)2SO4 as the nitrogen source, 10 mM potassium phosphate buffer) to a starting OD_600_ of 4. The cells were induced with 0.5% MeOH (v/v) and incubated at 25°C or 30°C and 280 rpm. For the following 48 hours the cells were fed twice per day with 2% MeOH (v/v). Samples for monitoring of cell growth (OD_600_) and metabolite concentrations were taken at the end of the cultivation and the best producing clones were identified and selected for further screening based on the calculated Y_P/X_.

### Shake flask cultivations

A screening of generated strains was performed in small scale shake flask cultivations. As in the 24-deep well plate screenings, a two-phase system was used [40]. For biomass production, strains were inoculated in a 10 mL YPG preculture (yeast extract 10 g·L^-1^, soy peptone 20 g·L^-1^, glycerol 20 g·L^-1^) overnight at 30°C and 180 rpm. Cells from the preculture were subsequently used for inoculation of the main culture with a starting OD_600_ of 4. For that, the cells were cultivated in 250 mL wide neck flasks, with a starting volume of 25 mL buffered YNB, at 30°C and 180 rpm. Cells were induced with 0.5% MeOH (v/v) and fed twice a day with 2% MeOH (v/v) for approximately 70 hours. Cell growth (OD_600_) was monitored throughout the cultivation and extracellular metabolite concentrations (MeOH, IA) were quantified by high-performance liquid chromatography (HPLC), as described below.

To investigate the tolerance to itaconic acid, a toxicity screening was performed in shake flasks as described above with minor changes; YNB medium was supplemented with IA to reach final concentrations of 25 g·L^-1^, 50 g·L^-1^, 100 g·L^-1^ respectively, pH was adjusted to 5.5 with KOH. Throughout the toxicity screening pH was measured and viability investigated via PI-staining as described below.

### Fed-batch cultivation

Fed batch cultivations were carried out in a DASGIP parallel bioreactor system (Eppendorf bioreactors SR0700ODLS and SR1000ODLS). To prepare the bioreactor inoculum, 100 mL YPG in a 1000 mL shake flask was inoculated with 1 mL of a working cell bank from the respective strain and incubated at 25°C, 180 rpm overnight (∼16 hours). Three hours before batch inoculation 50 mL YPG was added to each shake flask. Cells equivalent to reach a batch OD_600_ of 1 were used to inoculate the batch medium, modified BSM medium (13.5 mL·L^-1^ H_3_PO_4_ (85%), 0.5 g·L^-1^ CaCl_2_·2H_2_0, 7.5 g·L^-1^ MgSO_4_·7H_2_O, 9 g·L^-1^ K_2_SO_4_, 2g·L^-1^ KOH, 46.51 g·L^-1^ glycerol, 4.4 g·L^-1^ citric acid monohydrate, 0.25 g·L^-1^ NaCl, 8.70 mL·L^-1^ Biotin (0.1 g·L^-1^), 0.1 mL·L^-1^ Glanapon2000, 19.14 mL·L^-1^ 25% NH_3_, 4.35 mL·L^-1^ PTM0 (6.00 g·L^-1^ CuSO_4_⋅5H_2_O, 0.08 g·L^-1^ NaI, 3.36 g·L^-1^ MnSO_4_⋅ H_2_O, 0.20 g·L^-1^ Na_2_MoO_4_⋅2 H_2_O, 0.02 g·L^-1^ H_3_BO_3_, 0.82 g·L^-1^ CoCl_2_, 20.00 g·L^-1^ ZnCl_2_, 65.00 g·L^-1^ FeSO_4_⋅7 H_2_O, 5 mL·L^-1^ H_2_SO_4_ (95–98%)). Dissolved oxygen concentration was controlled at 30% by adjusting the stirrer speed, inlet gas flow and O_2_ concentration, where 200 rpm, 9.5 sL·h^-1^ and 21% were the minimal setpoints, 1250 rpm, 50 sL·h^-1^ and 100% were the maximum setpoints.

In the preliminary fermentation, the batch phase was followed by an 8 hour phase of glycerol-medium feed (550 g·L^-1^ 99% glycerol, 10.4 mL·L^-1^ PTM0, 10.4 mL·L^-1^ Biotin (0.2 g·L^-1^)) at a rate of 0.225·t+1.95, again followed by a cofeed phase of 17.5 hours where the glycerol-medium feed rate decreased over time (3.65-0.111·t) as the MeOH feed increased (0.028·t+0.6). After the cofeed phase the MeOH feed was continued at the same rate for the remaining time of the fermentation. The pH was maintained at 5.5 using 25% NH_3_ to counteract the acidification caused by the cultivation.

### Modification of fed-batch cultivations

To prevent continuous biomass accumulation in the remaining fermentations the base used for pH control was changed to 5 M KOH, and the feeding strategy was altered. The glycerol concentration of the modified mBSM was decreased to 33 g·L^-1^ to allow 20 g·L^-1^ of biomass at the end of the batch phases. The batch phases of ∼20 hours were followed by four hours of a limited glycerol feed phase (500 g·L^-1^ glycerol), with no additional trace elements or biotin. The limited glycerol feed phase was performed to reach a biomass of ∼40 g·L^-1^ and to induce the MeOH inducible genes [42]. A limited MeOH feed was then applied for the rest of the cultivation. The MeOH feed was adapted throughout the respective fermentation to prevent MeOH depletion as well as toxification.

### Sampling and productivity analysis of fed-batch cultivations

For all fed-batch cultivations samples were taken with time intervals for analysis of OD_600_, microscopy, dry cell weight (DCW) and HPLC, furthermore viability and samples for RNA extraction were included in some fermentations. For the respective fermentations, the bioreactor model, starting volume of the batch phase, final volume, feed-medium rate and MeOH feed rate are noted with the final biomass and IA-titers are noted in Table S2. Specific growth rates and productivity parameters were calculated according to the equations listed in Table 2.

**Table 2.**
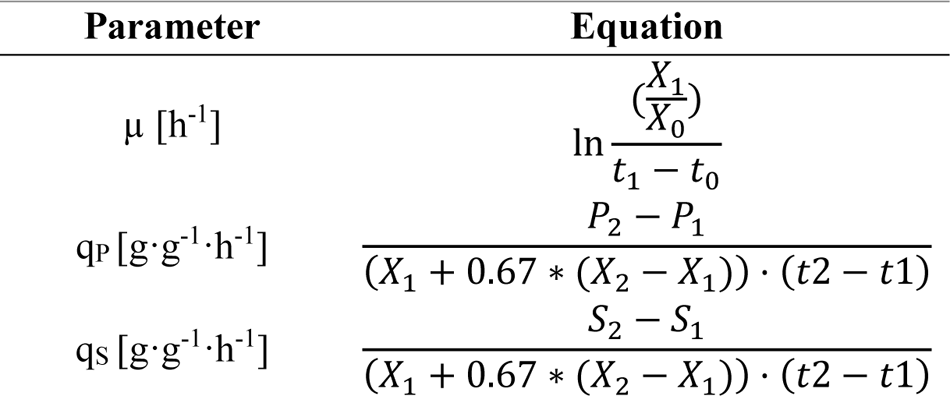
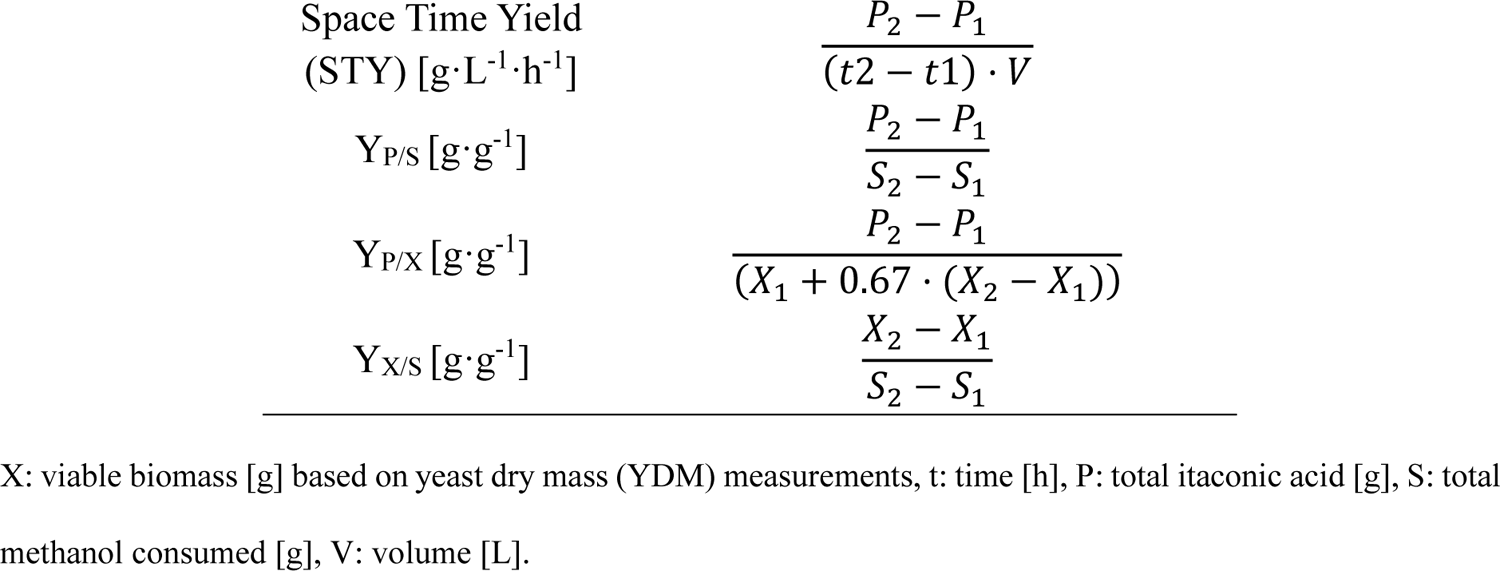
Equations used to calculate process parameters.

### HPLC Measurements

A Biorad Aminex HPX-87H HPLC column (300 × 7.8 mm) was used for the HPLC measurements. H_2_SO_4_ at a concentration of 4 mM was used as mobile phase, with a 0.6 mL·min^-1^ flow rate at 60°C. IA was measured with a photodiode array detector (SPD-M20A, Shimadzu) at 254 nm. MeOH and glycerol concentrations were quantified with a refraction index detector (RID-10A, Shimadzu). The chromatograms obtained with both detectors were used to confirm the absence of byproducts in relevant concentrations. Prior to HPLC measurement, samples were vortexed and centrifuged at full speed (16,100 g) for 5 min at room temperature. After centrifugation, the supernatant of each sample was mixed with H_2_SO_4_ (40 mM) resulting in a final concentration of 4 mM as the mobile phase. The supernatants were then filtered through a 0.22 μm filter into vials for the HPLC analysis.

### PI staining

To measure the viability of fed-batch yeast cultures, samples were diluted in PBS (0.24 g·L^-1^ KH_2_PO_4_, 1.8 g·L^-1^ Na_2_HPO_4_ · 2 H_2_O, 0.2 g·L^-1^ KCl, 8.0 g·L^-1^ NaCl) to an OD_600_ of 0.1-0.4 and stained with a propidium iodide (PI) stock solution to final concentrations of 2.0 M. PI acts as a DNA-intercalating fluorescence dye that permeates cells with compromised cell membranes thereby indicating cell death. When PI binds to DNA it leads to a strong increase in red fluorescence (excitation/emission at 480/630 nm) [43]. Samples were analysed on a CytoFlex S Flow Analyser 4L 13 (Color, Beckman Coulter), where the percentage of viable cells was estimated based on PI negativity.

### RNA extraction and subsequent cDNA conversion

Throughout the fermentations cell pellets were harvested and quenched with 1 mL TRI Reagent (SigmaAldrich) and frozen until further processing. RNA from 25 mg of yeast cell mass per 1 mL TRI Reagent was extracted by lysing the cells with ∼500 µL glass-beads at 5.5 m/s for 2·40s. Samples were then incubated at RT for five minutes before addition of 200 µL chloroform and vortexing for 15 seconds and incubation for 10 minutes at RT. To separate the RNA from DNA and protein, a centrifugation step was performed (4°C, 10 min, 10000 g) and the aqueous phase was mixed with 500 µL 2-propanol and incubated at RT for 8 min. The RNA was then pelleted by an additional centrifugation step (4°C, 10 min, 10000 g) and subsequently washed with 70% ethanol. When all ethanol had evaporated the RNA was solubilized with 80 µL water (65°C, 12 min). Residual genomic DNA was degraded with the DNA-free™-kit (Invitrogen) following the suppliers’ protocol.

The quality, purity and concentration of the isolated RNA was analysed with Nanodrop and visual analysis of an SYBR Safe agarose gel, and the absence of gDNA was verified by a PCR with primers homologous to the gene, *ACT1* and subsequent gel electrophoresis. cDNA synthesis was performed with the Biozym cDNA Synthesis Kit in accordance with the manufacturer’s protocol, using oligo d(T)23 VN (NEB) primers.

### Extraction of genomic DNA (gDNA)

Genomic DNA of the respective *K. phaffii* strains was extracted from overnight cultures using the Wizard Genomic DNA purification kit (Promega Corp., USA) according to the manufacturer’s instructions. The quality, purity and concentration of the isolated gDNA was analysed with Nanodrop.

### Analysis of relative gene copy numbers and gene expression

RT**-**qPCR was performed by mixing 2x qPCR S’Green BlueMix (Biozym Blue S’Green qPCR Kit), cDNA, water and qPCR primers (Table S1) in accordance with the supplier’s instructions. All samples were tested in technical duplicates and negative controls, without cDNA, were included for all primers. The quantitative gene expression analysis was performed using a real-time PCR cycler (Rotor Gene, Qiagen) and subsequent data analysis using the Comparative Quantitation (QC) method of the Rotor Gene software.

The GCNs of the *K. phaffii* multicopy strains (MC) were estimated by comparing the relative quantification of the interest sequences of the heterologous genes to that of the parent strain, cadA+mttA+mfsA_pGAP_, in which a single copy of the respective heterologous genes had been inserted locus-specifically via homologous recombination guided by CRISPR-Cas9. The GCNs were subsequently estimated relative to the corresponding parent strain using the threshold cycle (*ΔΔCT*) method [44].

The differences in gene expression amongst strains in different bioreactors were estimated by comparing the relative quantification of the respective reactor to the reference bioreactor at the same sampling time point. All signals were normalized to the constitutively expressed reference gene *ACT1* (PP7435_Chr3-0993).

## Results

### Enabling itaconic acid production from methanol in *K. phaffii*

Due to the absence of genes encoding a cis-aconitate decarboxylase, naturally occurring strains of *K. phaffii* are incapable of producing itaconic acid (IA). A recent study [45] has shown that a synthetic autotrophic *K. phaffii* could be engineered to produce IA from CO_2_ by insertion of a single gene, *cadA* from the natural producer *A. terreus*. The IA productivity was further improved by the insertion of another *A. terreus* gene encoding the mitochondrial cis-aconitate transporter, *mttA*.

In this study, a wild-type *K. phaffii* CBS7435 strain was engineered to produce IA from methanol (MeOH) by the integration of three heterologous genes from *A. terreus* in the *K. phaffii* genome. Initially, *cadA* encoding for the key enzyme was placed under the control of the strong MeOH-inducible alcohol oxidase 1 promoter (p*AOX1*) [38,46] resulting in the cadA strain. Next, *mttA* was inserted under the regulation of a moderately strong constitutive promoter (p*POR1*) [38], referred to hereafter as the cadA+mttA strain. Lastly, the *A. terreus* gene *mfsA*, encoding for a cytoplasmic membrane transporter specific to IA export, was integrated generating the cadA+mttA+mfsA strains. Previous studies have shown that modulating the transporter activity *in vivo* is crucial for optimal cell metabolism and productivity [45,47]. For that reason different promoters (p*FDH1*, pGAP, p*POR1*, p*FBA1* [38,46]) were tested for the expression of *mfsA*, in the cadA+mttA strain background. A 24-deep well plate screening was performed to evaluate the IA production of the newly generated cadA+mttA+mfsA strains, the IA yield, Y_P/X_, is given in Figure 1a. Notably, a strong promoter regulation of *mfsA* was beneficial for producing IA yielding comparable results obtained with the strong constitutive pGAP and the MeOH-inducible p*FDH1* promoter. The best-producing cadA+mttA+mfsA_pGAP_ strain was selected arbitrarily for further experiments, based on the screening results, and the benefits of having a constitutive expression of the product exporter.

**Figure 1:**
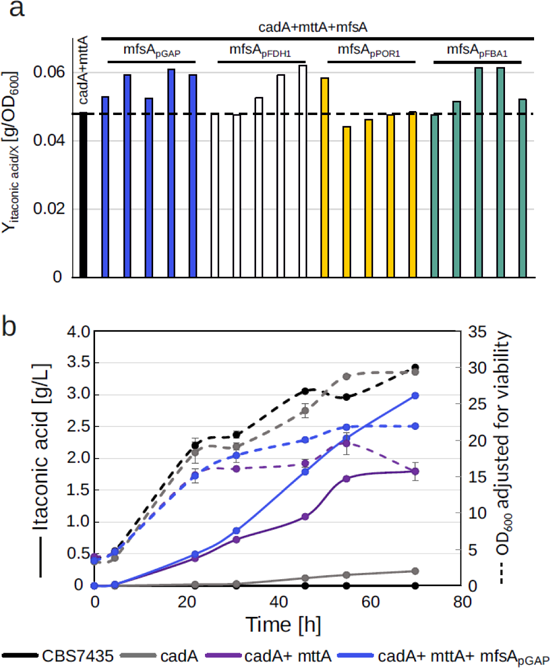
Fine-tuning promotor balance of the MfsA transporter is beneficial for itaconic acid production. **a)** Itaconic acid yields of cadA+mttA+mfsA strains after 48 hours of cultivation at 25°C for choice of best promoter for *mfsA* expression, **b)** Comparative shake flask cultivation of the CBS7435, cadA, cadA+mttA, cadA+mttA+mfsA_pGAP_ performed at 30°C. Throughout the cultivation samples were taken for OD_600_, HPLC and pH measurements, and for investigation of the viability of the strains with PI-staining. The OD_600_ is adjusted to exclude unviable cells. All strains were cultivated in duplicates, and average values and standard deviations for growth and itaconic acid production are shown.

To gain a comprehensive overview of how the expression of various genes of the IA pathway impacts productivity, we cultivated the strains cadA, cadA+ mttA, and cadA+mttA+mfsA_pGAP_ along with the CBS7435 strain as a reference in a comparative shake flask screening (Figure 1b), where samples for HPLC, pH measurements and viability (Figure S1) were taken at time intervals. While expression of *cadA* alone yielded 0.23 g·L^-1^ after 70 hours, additional expression of *mttA* improved the IA-titers eightfold to 1.8 g·L^-1^. Additional expression of *mfsA* further boosted the production with an additional 65% reaching 3.0 g·L^-1^ after 70 hours (Figure 1b). Intriguingly, growth was not affected by the expression of *cadA*, whereas additional expression of *mttA* reduced growth by 47% and whereas additional expression of *mfsA* restored the growth to 75% of the cadA strain.

### Itaconic acid production from methanol in fed-batch cultivations is feasible

To investigate whether the high IA productivity of the cadA+mttA+mfsA_pGAP_ strain was transferable to a larger scale, a fed-batch cultivation was performed with cadA+mttA+mfsA_pGAP_ alongside the cadA+mttA strain as a reference. The initial bioreactor setup for IA production was inspired by the setup for protein production of *K. phaffii* [48], since this was the standard for *K. phaffii* bioprocesses and as organic acid production from MeOH at that time had not been established in our laboratory. Therefore a standard four phase protocol was used, consisting of an initial glycerol batch phase for biomass production reaching 32 g·L^-1^ of biomass after 27 hours, a 8 hour glycerol-medium feed phase followed by a 17 hours long co-feed phase for a gradual MeOH gene activation [42], and finally a pure MeOH feed phase for organic acid production. During the MeOH feed phase the biomass increased from 85 g·L^-1^ to 140 g·L^-1^. The dissolved oxygen concentration (DO) was controlled at 30% and pH was maintained at 5.5 by an automatic addition of 25% NH_3_. In Figure 2 the IA and growth profiles of the fermentation are shown.

**Figure 2:**
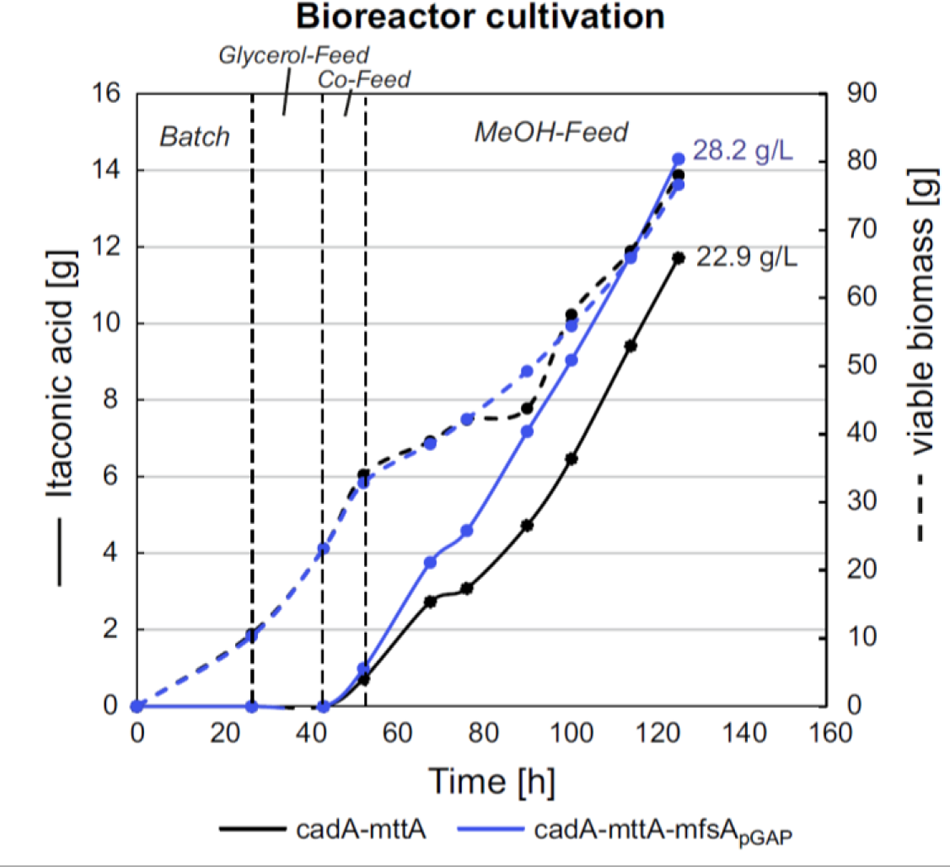
Upscaling of cadA+mttA+mfsA_pGAP_ cultivation leads to high itaconic acid production. Growth and production profiles are shown for the cadA-mtta and cadA+mttA+mfsA_pGAP_ strains. To ensure comparability of results despite varying cultivation volumes resulting from continuous MeOH and base feed, total amounts rather than concentrations are presented. The final IA-titers [g·L^-1^] are also shown. Time axis corresponds to the whole cultivation time including all four phases, i.e. batch, glycerol feed, co-feed and MeOH feed phase.

The 20% increase in productivity observed in the shake flask screening (Figure 1b) was reproducible on a larger scale, reaching 22.9 and 28.2 g·L^-1^ of IA by cadA+mttA and cadA+mtta+mfsA_pGAP_ respectively, resulting in an overall increase of 22% by the additional expression of *mfsA* (Figure 2) and a 40% increase in the yield on MeOH (Table 3). However, despite these improvements, the average yield per gram of MeOH remained at 0.1 g·g^-1^, primarily due to significant biomass accumulation (Table 3 and Table S2). This continuous biomass accumulation was expected given the fermentation strategy, which was originally developed for protein production favoring high cell densities. The continuous growth was enabled by a nutrient-rich complex medium (modified mBSM) and an ongoing nitrogen replenishment through the addition of NH_3_ as a base control.

**Table 3:**
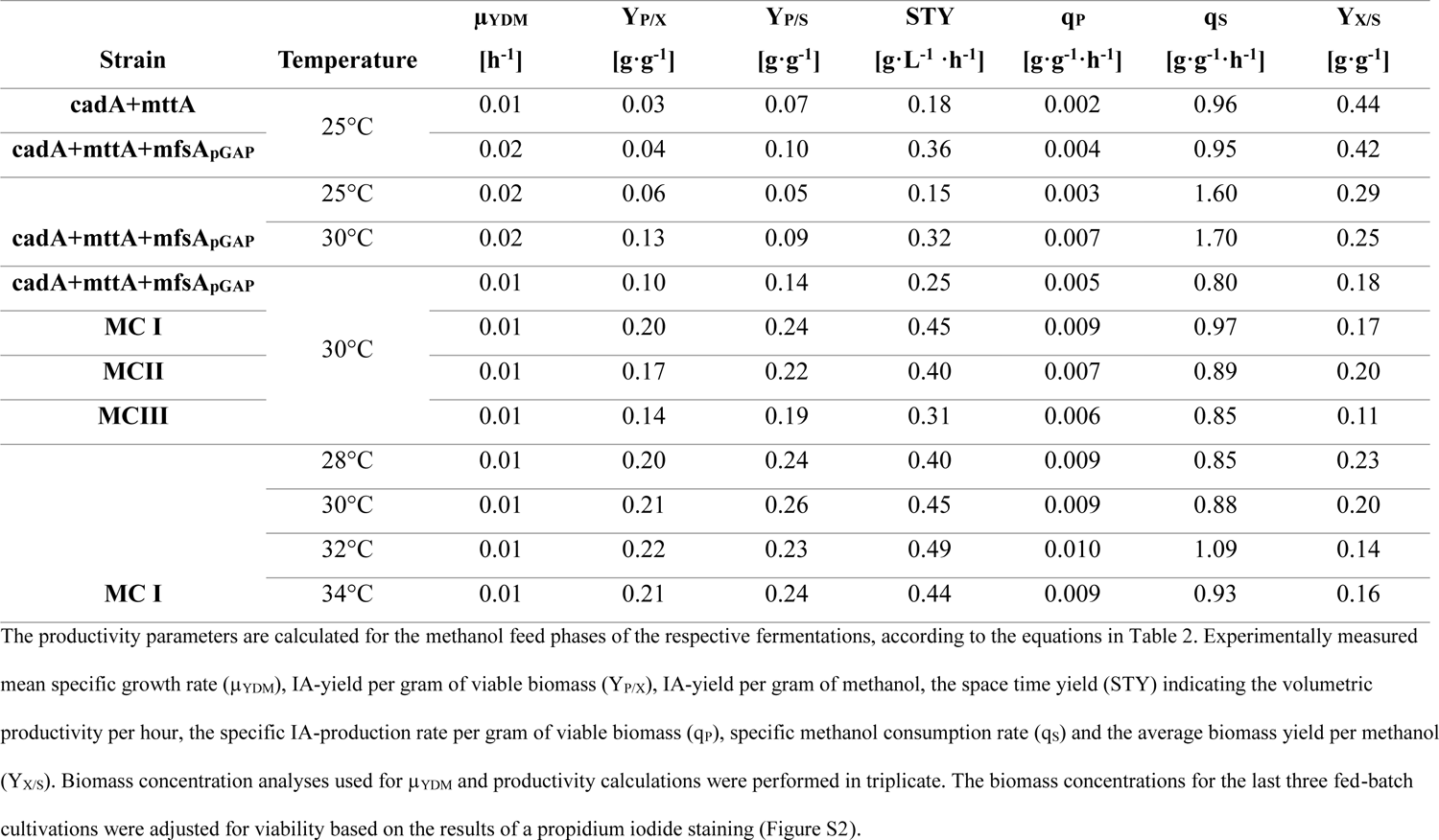
Average values of key productivity parameters obtained from the methanol feed phase of fed-batch cultivations.

### Increasing fermentation temperature by 5°C doubles itaconic acid yields

To further optimize the IA production, the following adjustments of the fermentation process were applied. First, to prevent high biomass formation as observed in the previous fermentation a nitrogen-limitation was introduced by changing the base from the 25% NH_3_ to 5 M KOH. Consequently, only nitrogen present in the batch medium was utilized by the cells for biomass formation. Second, the glycerol-feed and co-feed phases were replaced by a restricted glycerol feed lasting 4 hours. These adjustments made the process less time-consuming and resulted in a comparable effective activation of the MeOH-utilizing genes [42]. And finally, the process optimization temperature was elevated from 25°C to 30°C. The IA production and growth profiles of the fermentation are shown in Figure 3. Throughout the fermentation samples were taken and analysed as in the previous fed-batch cultivation with additional samples for PI-staining to gain a better insight of the cell viability throughout the process (Figure S2a). As a clear trend was already observed after 64 hours, the fermentation was stopped and evaluated.

**Figure 3:**
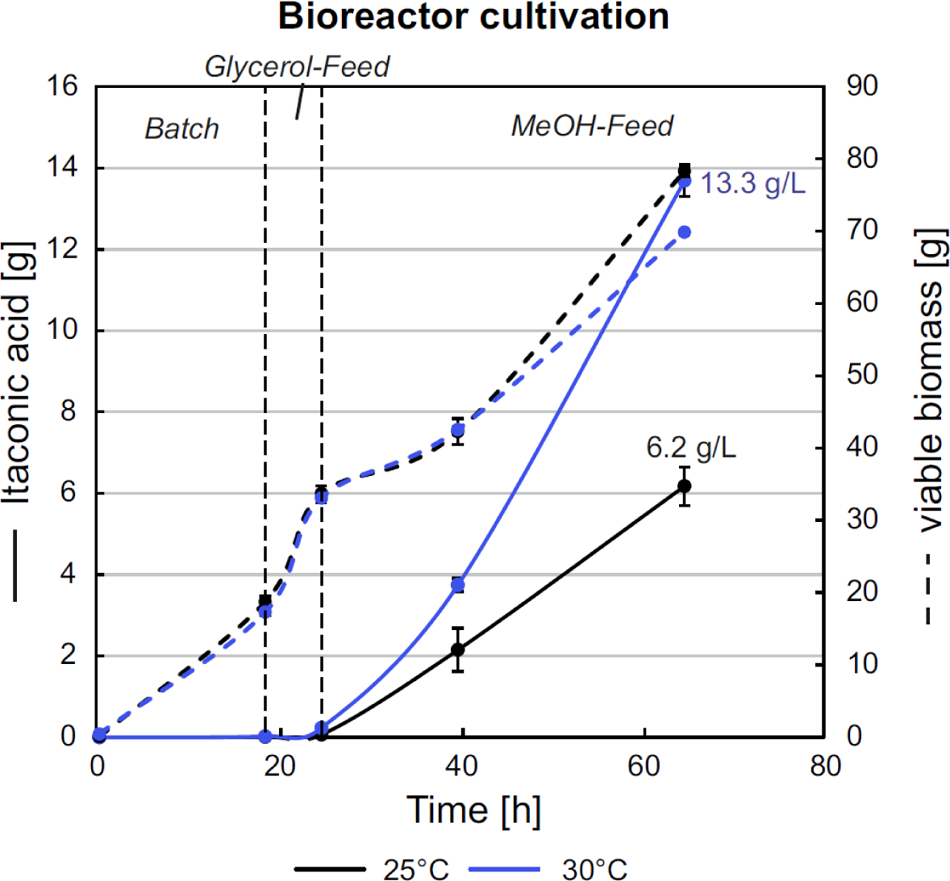
Temperature dependency of itaconic acid production in *K. phaffii* favors a higher temperature. Growth and production profiles are shown for the cadA+mttA+mfsA_pGAP_ strain cultivated at 25°C and 30°C. To ensure comparability of results total amounts rather than concentrations are presented. The final IA-titers [g·L^-1^] are also shown. The time axis corresponds to the whole cultivation time including all three phases, i.e. batch, glycerol feed and MeOH feed phase. The growth profile at 30°C is markedly lower than at 25°C, which in combination with the improved IA-productivity clarifies that a cultivation temperature of 30°C is favourable.

Intriguingly, the production efficiency increased markedly along with the elevated process temperature. As indicated in Figure 3 and the calculated productivity parameters in Table 3, the average yield per biomass was more than doubled (+123%), whilst the conversion of MeOH into IA was doubled simultaneously with a 13% decrease in the biomass accumulation after only 64 h. In terms of titers, elevation of the process temperature by 5°C pushed the IA concentrations from 6.2 g·L^-1^ to 13.3 g·L^-1^ after only 40 hours of MeOH feed phase, suggesting that a higher process temperature is significantly beneficial for IA production.

#### Integrating multiple copies of itaconic acid pathway genes improves production when the right balance between genes is achieved

Elevating the gene copy number (GCN) of heterologous biosynthesis genes is a common strategy to enhance productivity of strains. In other synthetic IA producers it has recently been demonstrated that increasing the GCN of *cadA* improves IA-productivity [49,50]. Similarly, the push-and-pull effect has previously been shown to promote production of other organic compounds [45,51,52]. Hence, we aimed to increase the GCN of *cadA*, *mttA* and *mfsA* either individually or in a combined approach. For that the cadA+mttA+mfsA_pGAP_ strain was transformed with plasmids for multicopy integration of either *cadA*, *mttA* or *mfsA,* or the following combinations: *cadA*+*mfsa* and *cadA*+*mttA*+*mfsA*. A set of the generated multicopy clones (MC) was cultivated in 24-deep well plates for 48 hours (Figure 4). Additionally, all clones were tested for their GCN via a quantitative PCR approach (Figure S3a-c). Based on the OD_600_ and the IA concentration, the productivity of the individual clones was evaluated by calculating the IA-yield per biomass (Figure 4).

**Figure 4:**
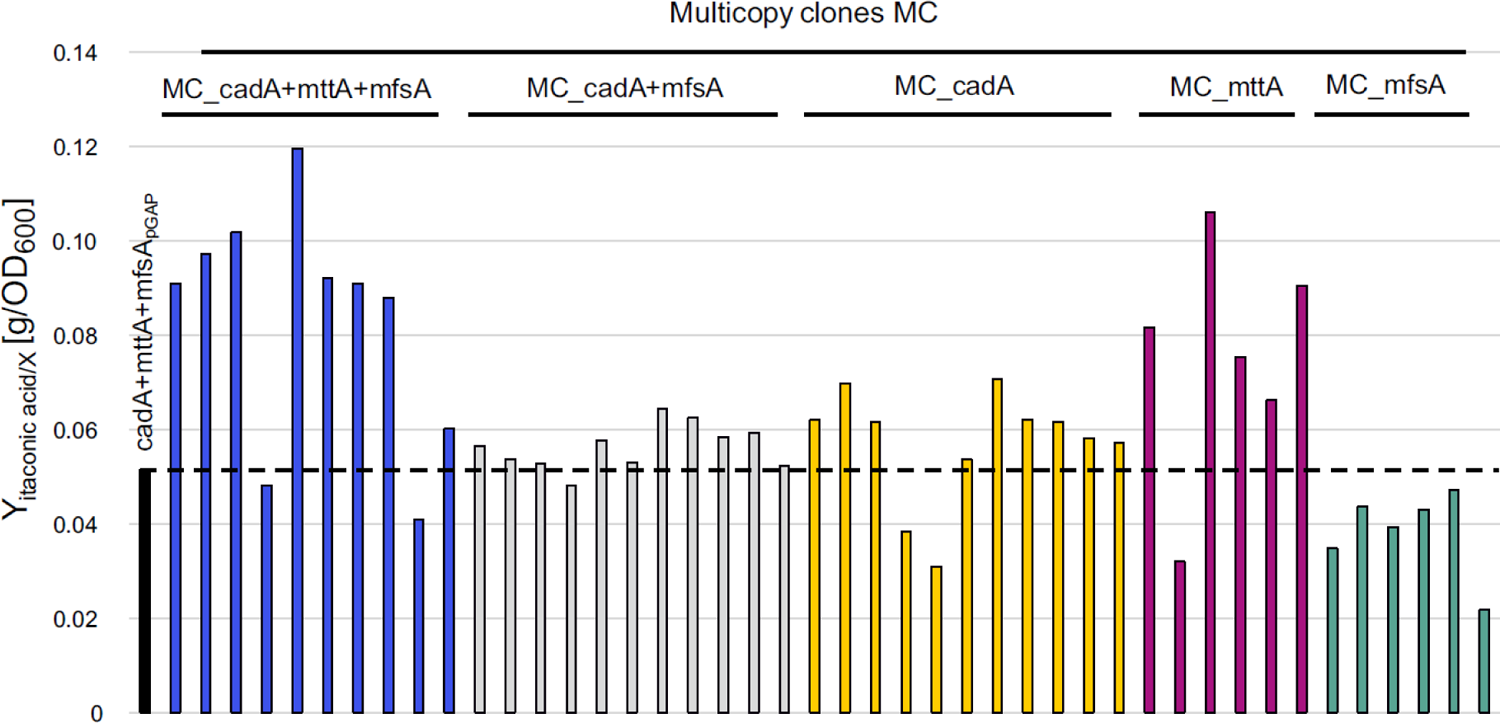
Integration of multiple copies of the heterologous genes doubles itaconic acid yield. The yield per biomass (OD_600_) was evaluated for the individual multicopy strains (MC) and compared with the cadA+mttA+mfsA_pGAP_ parental strain after 48 hours of cultivation with a 2% MeOH feed in 24-deep well plates.

Upon analysing the IA productivity alongside the gene copy numbers (GCN) of the heterologous genes (Figure 4 & Figure S3a-c), it is evident that inserting multiple copies of *cadA* or *mfsA* as individual genes, as well as the multicopy combination cadA+mfsA, has a limited or even negative effect on IA production in most clones. The majority of the MC_cadA+mttA+mfsA clones showed increased yields, indicating that multicopy integration of all three heterologous genes, *cadA*+*mttA*+*mfsA*, is necessary to improve the IA yield, provided that the optimal balance of the three genes is reached. Surprisingly, several of the MC_mttA clones showed a high productivity per biomass, however, the production titers were low, and so the high yields were only enabled by a low biomass accumulation (Figure S3d). When analysing the OD_600_ with the GCN, a correlation between a high GCN of *mttA* with an impaired growth in both the MC_mttA and the MC_cadA+mttA+mfsA clones is observed. Despite the high Y_P/X_ the strains with a high *mttA* GCN were likely unfit for longer fed-batch cultivations.

Based on the 24-deep well plate screening (Figure 4), and subsequent shake flask cultivations (Figure S4), three clones (hereafter referred to MC I, MC II and MC III) were selected for a fed-batch cultivation along with the cadA+mttA+mfsA_pGAP_ parent strain. MC I and II were chosen as they had shown the highest IA productivity alongside parent-like growth, whilst MC III was selected for its relatively high productivity combined with a decreased growth phenotype (Figure S3). The fed-batch cultivation was performed as the previously described three-phase cultivation at 30°C. Samples were extracted continuously to allow for analysis of growth, IA production and viability, furthermore samples for RNA extraction were included.

Going in line with the results from the previous screenings, all MC strains showed improved IA productivity (Figure 5a). Specifically, MC I showed the highest IA titers of 49.5 g·L^-1^, followed by MC II with 45.6 g·L^-1^ and MC III with 32.9 g·L^-1^. The parent strain cadA+mttA+mfsA_pGAP_ reached only 27.2 g·L^-1^. MC III exhibited a notable reduction in biomass formation while growth rates remained similar between the parent strain, MC I, and MC II (Figure 5a). The MC III shows a decline in biomass after 80 hours of cultivation, this is in line with the previous experiments and an observed correlation between growth impairments and high *mttA* GCN. The *mttA* GCN of MC III is double that of the other strains (Figure 5b). High GCN of *cadA* and *mfsA*, in combination with a slight increase in the GCN of *mttA*, are beneficial for IA production, as implicated by MC I and MC II (Figure 5a-b). To verify if a higher GCN correlates with an elevated expression of the genes, RNA was extracted at different time points throughout the fermentation and subsequently cDNA was analysed via RT-qPCR. As gene expression was found to be highest at the end of fermentation, this particular time point is depicted in Figure 5c. In alignment with the GCN, expression levels of the heterologous genes *cadA* and *mttA* were elevated accordingly, suggesting a direct correlation between GCN and gene expression and indicating that the multiple copies are active. Interestingly, the trend between GCN and fold change is not observed for *mfsA* expression in all three MC clones. Instead, the MC III, with a high *mttA* GCN and an extremely high *mttA* expression level, shows the highest *mfsA* expression, suggesting that the high *mttA* expression affects the expression of *mfsA*. Overall, with generating multicopy strains IA productivity could markedly be improved, resulting in an improvement of the average STY by up to 80% and Y_P/S_ up to 71% when compared to the parental strain under identical cultivation conditions (Table 3). The production efficiency of MC I was moderately higher than MC II, but when taking the observed viability drop for MC II (Figure S2b) into account, MC I did not only show the highest productivity, it also showed the highest robustness, making it the optimal strain for subsequent experiments.

**Figure 5:**
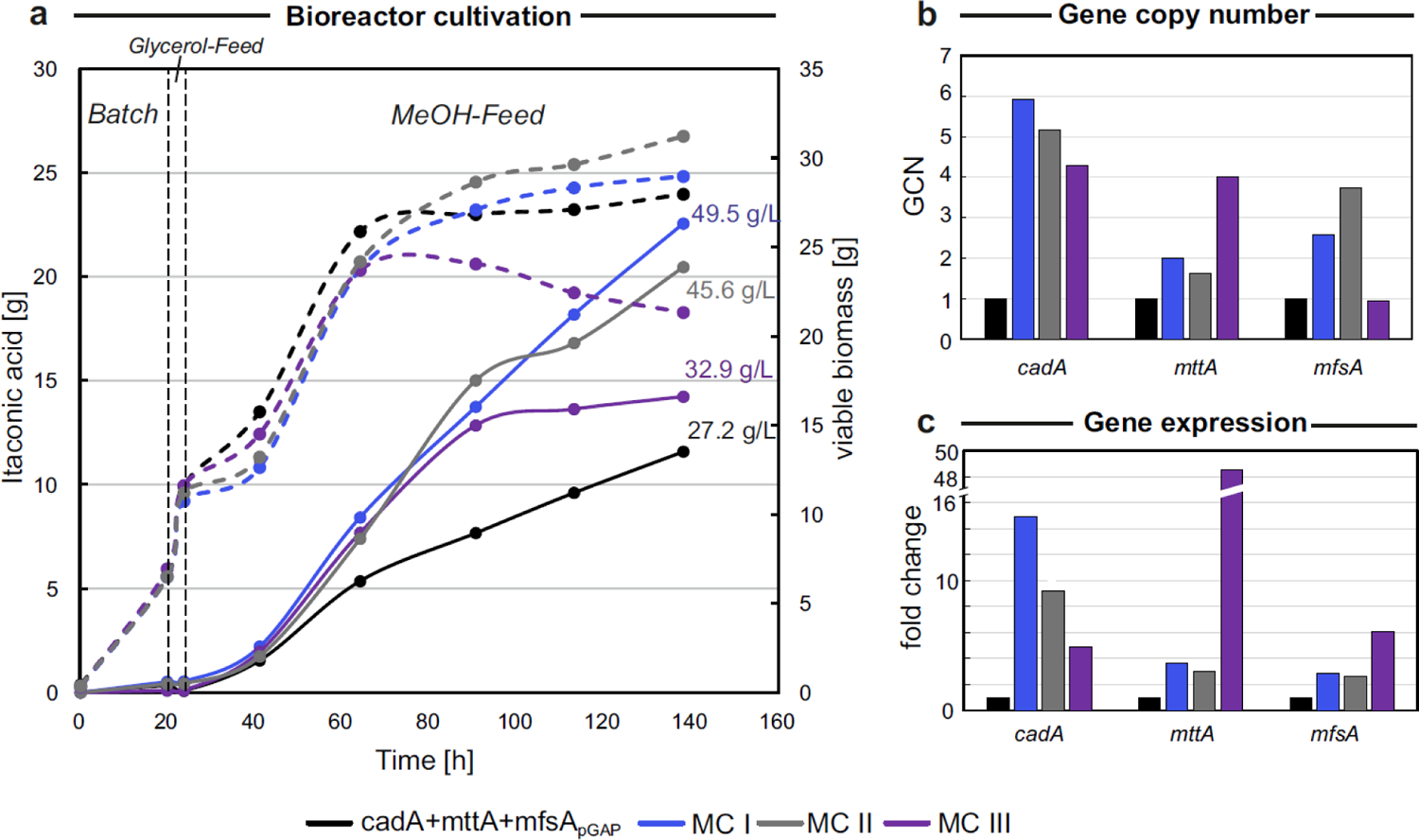
Balancing the gene copy number of the three heterologous genes enables titers of 49.5 g·L^-1^ after five days of fed-batch cultivation. **a)** Growth and production profiles are shown for the cadA+mttA+mfsA_pGAP_ strain and three selected multicopy strains (MC I, MC II and MC III) cultivated at 30°C. To ensure comparability of results total amounts rather than concentrations are presented, the final IA-titers [g·L^-1^] are also shown. The time axis corresponds to the whole cultivation time including all three phases, i.e. batch, glycerol feed and MeOH feed phase. All MC strains show an increased IA production compared to the parental strain. **(b)** Gene copy numbers of the three multicopy strains relative to the parental strain with single copy integration of all three heterologous genes. **(c)** The relative expression of the three heterologous genes at the fermentation for the three multicopy strains normalized to the expression of the parental strain at the same time point.

### Expression of *mfsA* improves tolerance to itaconic acid

High IA concentrations both within and outside the cells might burden the cells. Especially with high *mttA* expression, which increases the substrate availability for CadA, IA accumulation in the cytosol is probable. High intracellular IA concentrations are likely to have a negative effect on cell metabolism. Strong expression of *mfsA* can therefore be beneficial for the well-being of the cell, as it facilitates the export of IA and maintains a low cytosolic pool of the organic acid. To evaluate if (i) the generated strains are suitable for high IA production in general and (ii) differences exist in IA tolerance between the engineered strains, a toxicity screening was performed. Therefore, the cadA, cadA+mttA, cadA+mttA+mfsA_pGAP_, and the MC I strain were cultivated alongside the CBS7435 strain in presence of high external IA concentrations in the range of 25, 50 and 100 g·L^-1^ at a pH range of 5.5-5.7. OD_600_ and viability were measured throughout the screening and are depicted in Figure 6.

**Figure 6:**
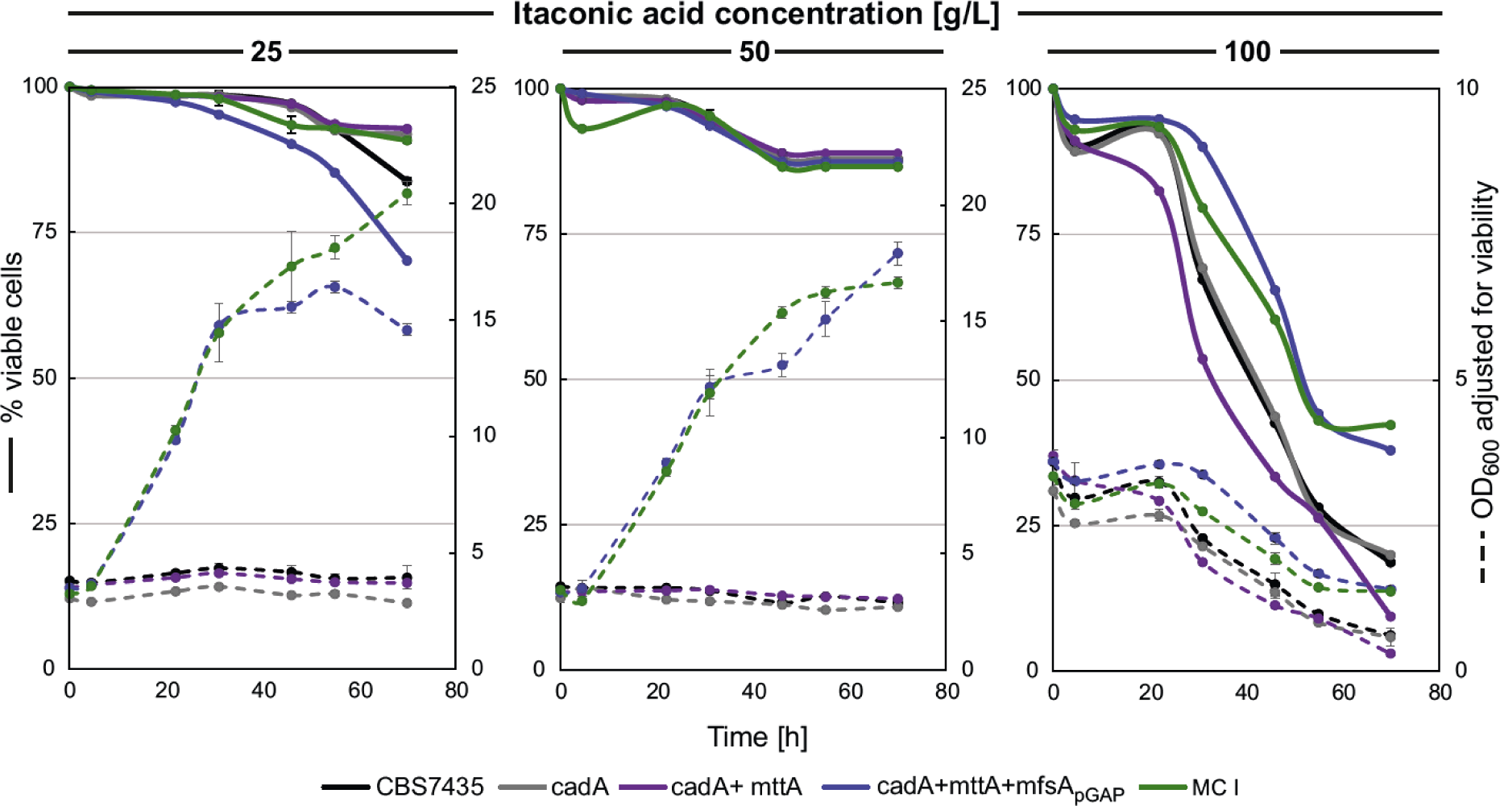
Expression of *mfsA* improves *K. phaffii’s* tolerance to extracellular itaconic acid. The four engineered strains were cultivated in parallel with CBS7435 in cultivations with high IA-concentrations (25, 50 and 100 g·L^-1^) at 30°C.

Interestingly, IA concentrations in the range of 25 g·L^-1^ inhibited growth of the CBS7435, cadA and cadA+mttA strain. Under these conditions, only strains expressing the MfsA transporter i.e., cadA+mttA+mfsA_pGAP_ and MC I, demonstrated growth. Similar results were obtained in cultivations with 50 g·L^-1^ of IA. Throughout these conditions, viability remained relatively stable, not dropping below 70%. However, at IA levels of 100 g·L^-1^, both the cadA+mttA+mfsA_pGAP_ and MC I strain were adversely affected in the utilized cultivation conditions, as observed in the declining biomass profile (Figure 6). Nonetheless, both strains exhibited better performance with their viability stabilizing at 38-42% compared to CBS7435, cadA, and cadA+mttA strains where viability dropped below 20%. These results clearly indicate that the expression of *mfsA* improves the tolerance to IA.

### Optimal process temperatures are between 30-32°C for the MC I strain in fed-batch bioreactor cultivations

Having successfully engineered a high IA producer strain, MC I, through multicopy gene integration, we proceeded with a final fermentation experiment aimed at optimizing the process temperature. For that the same three-phase fed-batch strategy was used, testing fermentation temperatures of 28, 30, 32 and 34°C. Samples were extracted for growth- and HPLC as well as for viability analysis. IA and growth profiles are depicted in Figure 7. The best results were observed within the temperature range of 30-32°C. While 32°C showed slightly better overall process parameters for production (Y_P/X_, STY, q_P_), the MeOH conversion to IA was more favourable at 30°C (Table 3). At 28°C biomass formation was improved rather than IA production, while 34°C resulted in a lower biomass and IA profiles. In all cases, strains maintained a high viability throughout the fermentation process (Figure S2c), at 34°C the viability did however drop to 70.5%. Overall, at temperatures between 30-32°C MC I was able to produce between 50-55 g·L^-1^ IA.

**Figure 7:**
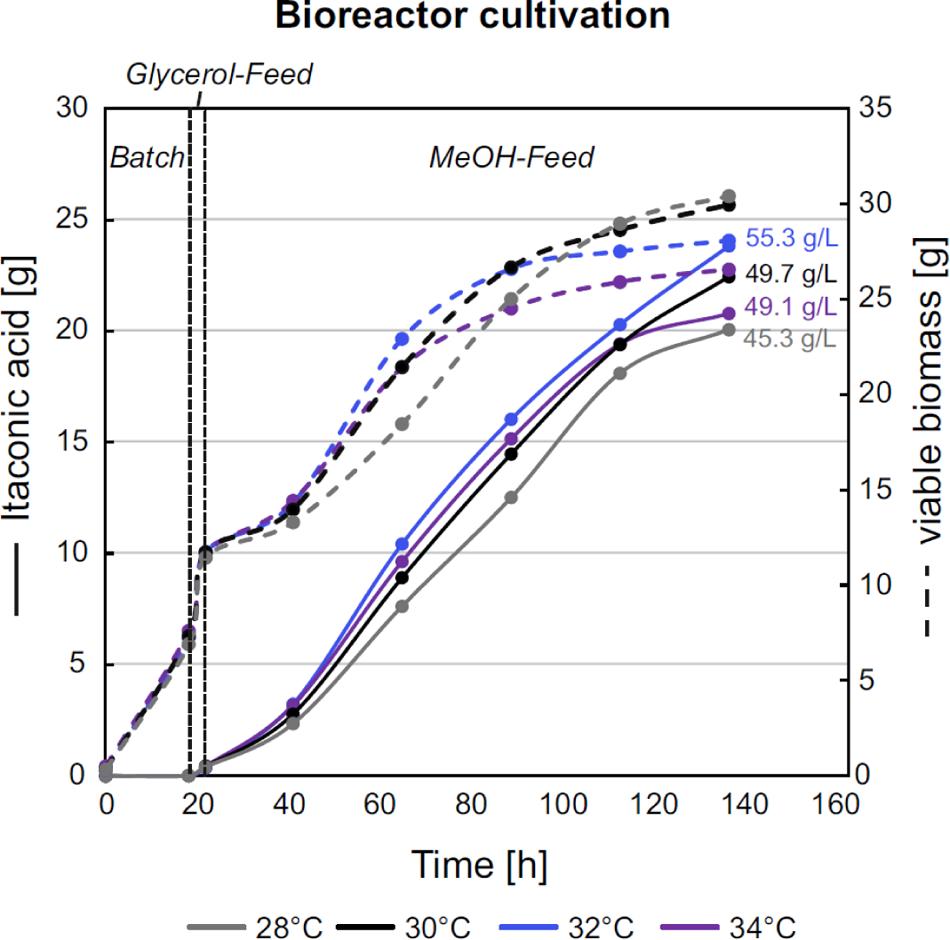
Fed-batch cultivation of the engineered *K. phaffii* strain shows highest productivity between 30 to 32°C. Growth and production profiles are shown for the MC I strain at different cultivation temperatures (28, 30, 32, 34°C). To ensure comparability of results total amounts rather than concentrations are presented, together with the final IA-titers [g·L^-1^]. Time axis corresponds to the whole cultivation time including all three phases, i.e. batch, glycerol feed and MeOH feed phase.

## Discussion

Traditionally, itaconic acid (IA) production has been primarily associated with filamentous fungi such as *Aspergillus spp.* or *Ustilago maydis* using glucose as a carbon source [53, 54–56]. However, the search for alternative hosts has been ongoing, driven by the need for more sustainable production methods and the exploration of diverse substrates. In this study we show that *K. phaffii* can produce IA from methanol (MeOH) reaching titers up to 55 g·L^-1^. This was achieved by a combinatorial approach focusing on strain engineering and process optimization.

In several natural non-producing organisms overexpression of *cadA* has enabled IA production from different carbon substrates, including C1 feed stocks [56]. Just to mention a few, *cadA* expression enabled *Methylorubrum extorquens* AM1 IA production from MeOH reaching titers of 31.6 mg·L^-1^[53], and autotrophic IA production from CO_2_ was enabled in strains such as *Synechocystis* sp. PCC6803 and a synthetic autotrophic *K. phaffii* strain, yielding 14.5 mg·L^-1^ and 0.53 g·L^-1^, respectively [45,57]. Also in our hands, expression of *cadA* enabled IA production in the range of 0.23 g·L^-1^ on MeOH in *K. phaffii.* The production capacity was however boosted with the coexpression of the MttA transporter, reaching titers of 1.8 g·L^-1^. Expression of *mttA* creates possibly a pull effect of cis-aconitate from the mitochondria to the cytosol, thereby providing more precursor molecules for conversion by CadA to IA [58,59]. Further, we demonstrated that by incorporating an *A. terreus* gene encoding a plasma membrane transporter, *mfsA*, under the control of a strong promoter, IA titers of 3.0 g·L^-1^ were reached in shake flask cultivations. The pivotal role of both transporters has previously been underscored by [60], a finding corroborated by our observations. Furthermore, our results suggested that utilizing a strong promoter for *mfsA* proves advantageous for IA production. Subsequent toxicity screenings revealed that MfsA expression also improved growth and viability of *K. phaffii* at high IA concentrations. This phenomenon is likely attributed to the efficient export of IA by the cell, thereby mitigating weak organic acid induced stress within the cellular environment.

Upscaling the process to bioreactor cultivations was successful, reaching titers of 28.2 g·L^-1^ with the strain expressing a single copy of all three heterologous genes *cadA*, *mttA* and *mfsA*. However, excessive biomass accumulation was a significant side effect, hence process optimization was required. The combination of nitrogen limitation and an increase in process temperature from 25 to 30°C resulted in a twofold increase in the MeOH yield for the cadA+mttA+mfsA_pGAP_ strain. Additionally, it led to a 13% reduction in biomass accumulation, favoring the conversion of MeOH to IA over biomass production. A substantial increase in productivity was anticipated, considering the prior evaluation of CadA activity [35]. Here, in vitro studies revealed a positive correlation between enzyme activity and temperature, peaking at 45°C in *A. terreus.* Further process optimization of the MC strains, exploring temperatures ranging from 28 to 34°C, revealed that the engineered strain exhibited optimal performance within the temperature range of 30-32°C. Notably, cellular metabolism was adversely impacted by 34°C leading to decreased viability over time and less efficient MeOH utilization compared to the temperature range of 30-32°C. This decline in cell viability is particularly undesirable for downstream processing of the fermentation broth, as reduced viability results in increased cell lysis and thus presence of unwanted cell debris [61].

Next to this, we could show that increasing the number of all heterologous genes markedly improved the potential of *K. phaffii* as an IA production host, as it increased IA productivity (0.49 g·L^-1^·h^-1^) simultaneously with an efficient conversion of MeOH into IA (0.26 g·g^-1^). In several organisms [49,50], the integration of multiple copies of *cadA* has proven beneficial for production. The instability of CadA was previously observed [31,62], though whether the necessity for additional cadA expression is linked to this instability remains unanswered at this point. However, our findings suggest that the integration of additional *cadA* copies had a minimal impact on IA production, unless it coincides with the multicopy integration of *mttA* and *mfsA*. The heightened expression of these two transport proteins enhanced the pull effect. Notably, strains harboring a high *mttA*-GCN showed severe growth impairments in comparison to the remaining multicopy strains. Similar results were obtained by [45], indicating that a strong promoter regulation for mttA expression severely impacts growth in the autotrophic *K. phaffi*i strain. A high MttA activity might efficiently drain the TCA cycle from its intermediate and result in accumulation of IA in the cytosol, thus possibly inflicting acid-induced stress within the cell. The additional expression of *mfsA* in the cadA+mttA+mfsA_pGAP_ and MC_cadA+mttA+mfsA strains possibly compensates for this phenotype as IA-toxification becomes less likely due to the increased export.

## Conclusion

Overall, the success of our IA production process using *K. phaffii* as a platform is attributed to the combined efforts of metabolic engineering and process engineering, emphasizing the importance of both aspects in developing efficient and sustainable production processes. Although our strain cannot yet compete with industrially relevant yields of 0.58 g·g ^-1^ produced by *A. terreus* [33] in terms of carbon conversion, one must consider the origin of the carbon, as the use of glucose in the current industrial process is unfavourable for a sustainable development of biobased production. Our process establishes for the first time a high-yield MeOH-based production system with a high IA-production rate of 0.49 g·L^-1^·h^-1^. This high production rate in combination with the advantages of MeOH as carbon source, the additional safety of *K. phaffii* as a non-pathogenic, well established production host and the minimal by-product formation streamlining downstream purification, highlight the relevance of our process. With additional metabolic and process engineering advancements, *K. phaffii* is poised to emerge as a significant industrial production host for IA utilizing MeOH.

## Supplementary Information

The supplementary material consists of one pdf file including Table S1 and S2 as well as Figure S1-S4.

## Acknowledgements

We want to thank Viktoria Kowarz and Bernhard Schmelzer for their great help in performing some of the fed-batch cultivations. M.M.S. was further supported by the Bioprocess Engineering Doctoral School of BOKU University.

## Author contributions

M.M.S., S.B., O.A. and D.M. contributed to the design of the work. M.M.S, V.M. and S.B. were involved in data acquisition. M.M.S., S.B., O.A. and D.M. were involved in data analysis. M.M.S., S.B. and D.M. wrote the manuscript. All authors read and approved the final manuscript.

## Availability of data and materials

The data that supports the findings of this study are included in this article /supplementary files. Further datasets used and/or analysed during the current study are available from the corresponding author on reasonable request.

## Declarations

### Ethics approval and consent to participate

Not applicable

### Consent for publication

Not applicable

