## Supplementary File for "Efficient production of itaconic acid from the single carbon substrate methanol with engineered *Komagataella phaffii*"

**Table S1**. **Primer names and sequences.**

| Name | Sequence | Comments |
|---|---|---|
| **Primers used for diagnostic verification of generated strains** | | |
| dia_GUT1_check_fwd | GATTCCAAACTGCAGGAACGCAG | Primers bind outside the GUT1 locus and are used for verification of GUT1 insertions. |
| dia_GUT1_check_rev | GAGGAGTCGGCAAAGTACCC | |
| dia_RGI_check_fwd | ATCAAACTTTTTGAATGGA | Primers bind outside the RGI locus and are used for verification of RGI-directed inserts. |
| dia_RGI_check_rev | CTATAAGAAACTGGAGACT | |
| BB2_check_fwd | CTGCGTTATCCCCTGATTCT | Primers used to verify integration of transcription unit in BB2 plasmids. |
| BB2_check_rev | GGGTGAGCAAAAACAGGAAG | |
| BB3_check_fwd | TTAGTATGCTGTGCTTGGGTG | Primers used to verify multicopy integration of the heterologous genes. The respective genes were told apart based on size observed via gel electrophoresis. |
| BB3_check_rev | GAGGTATGTAGGCGGTGCTA | |
| **Primers used for RT-qPCR analysis** | | |
| q_act1_fw | CCTGAGGCTTTGTTCCACCCATCT | Primers used to analyze gene expression via RT-qPCR |
| q_act1_rv | GGAACATAGTAGTACCACCGGACATAACGA | |
| q_cadA_fw | GGGTCGCGTGAGGATTGAGTTC | |
| q_cadA_rv | CTACCCGCAAGGGTTCGGTAT | |
| q_mttA_fw | GCAACACCTTCAACTGCGTCAA | |
| q_mttA_rv | GACCTTCTCGTAAACGGGGAACAT | |
| q_mfsA_fw | AGTGCGTCATCACCTTCGTC | |
| q_mfsA_rv | AGGGAGGAGTAGCCCATAGC | |

Primers used for diagnostic verification of the generated strains, for RT-qPCR analysis and for cDNA synthesis.

**Table S2. Process parameters of the respective fed-batch cultivations.**

| Total cultivation time [h] | MeOH phase time [h] | Strain | Temperature | Initial batch volume [mL] (Bioreactor model) | MeOH flow rate [$g \cdot L^{-1} \cdot h^{-1}$] | Feed-medium flow rate [$mL \cdot h^{-1}$] | Volume [mL] | Itaconic acid [g/L] | YDM [$g \cdot L^{-1}$] |
|---|---|---|---|---|---|---|---|---|---|
| 126 | 73 | cadA+mttA | 25 °C | 320 (SR0700ODLS) | $0.028 \cdot t+0.6$ | $0.225 \cdot t+1.95$ | 556.7 | 23.0 | 140.2 |
| | | cadA+mttA+mfsA$_{pGAP}$ | | | | $3.65-0.111 \cdot t$ | 552.2 | 28.2 | 138.8 |
| 65 | 40 | cadA+mttA+mfsA$_{pGAP}$ | 25 °C | 820 (SR1000ODLS) | 3.1 | 10.3 | 1028.5 | 6.2 | 76.1 |
| | | | 30 °C | | 3.1 | | 1053.3 | 13.3 | 66.3 |
| 138 | 114 | cadA+mttA+mfsA$_{pGAP}$ | 30 °C | 320 (SR0700ODLS) | 2.0 | 4 | 447.2 | 27.2 | 66.6 |
| | | MC I | | | 2.0 | | 476.1 | 49.5 | 62.6 |
| | | MCII | | | 1.9 | | 470.7 | 45.6 | 67.0 |
| | | MCIII | | | 2.5 | | 449.1 | 32.9 | 48.0 |
| 137 | 115 | MC I | 28 °C | 320 (SR0700ODLS) | 1.6 | 4 | 464.5 | 45.3 | 65.8 |
| | | | 30 °C | | 1.8 | | 472.3 | 49.7 | 63.7 |
| | | | 32 °C | | 2.2 | | 451.9 | 55.4 | 65.9 |
| | | | 34 °C | | 1.8 | | 443.2 | 49.1 | 60.6 |

Total cultivation time [h], duration of MeOH phase [h], strain, temperature, batch volume and reactor model, average MeOH flow rate [$g \cdot L^{-1} \cdot h^{-1}$], glycerol-feed flow rate [$mL \cdot h^{-1}$] and final volume [mL], final itaconic acid titer [$g \cdot L^{-1}$] and yeast dry cell mass (YDM) [$g \cdot L^{-1}$] are given.

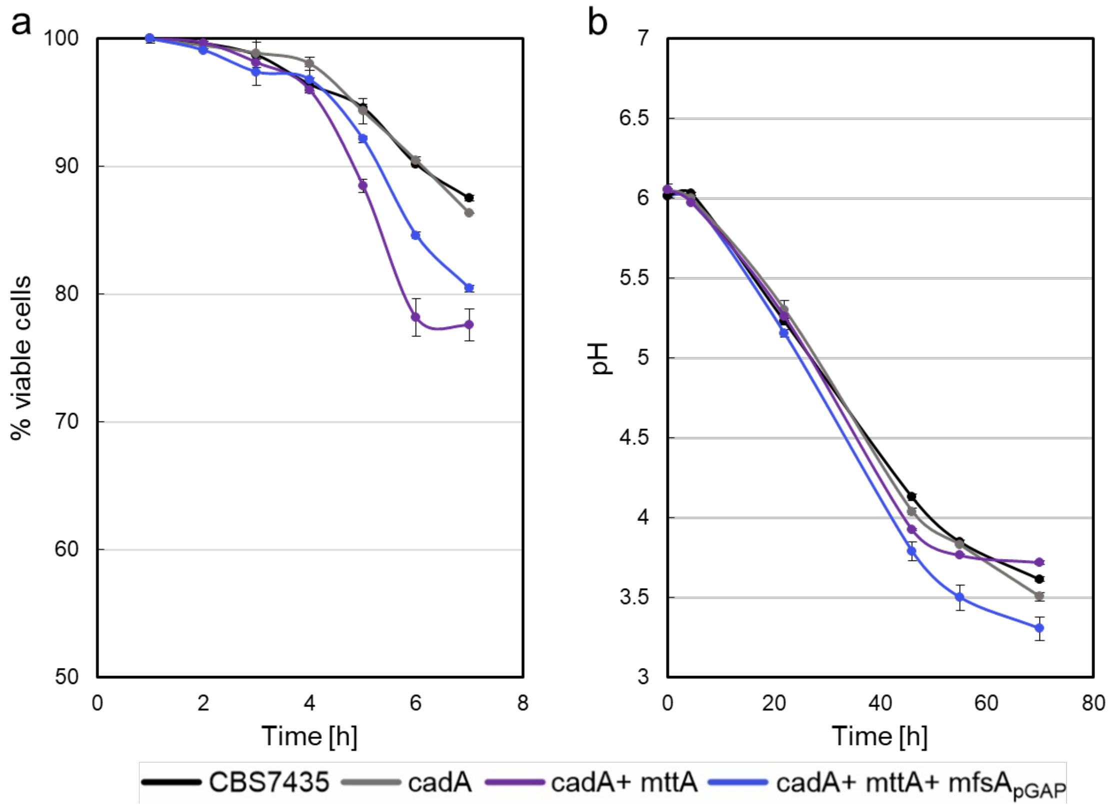


**Figure S1. Viability of strains during pH change.** During the comparative shake flask cultivation of the *K. phaffii* strain CBS7435 with the generated strains (cadA, cadA+mttA and cadA+mttA+mfsA$_{pGAP}$ **a)** viability of the strains was estimated with PI-staining at every sampling point, whilst **b)** pH was also measured. The strains were cultivated in biological duplicates and averages and standard deviations are shown for both viability and pH.

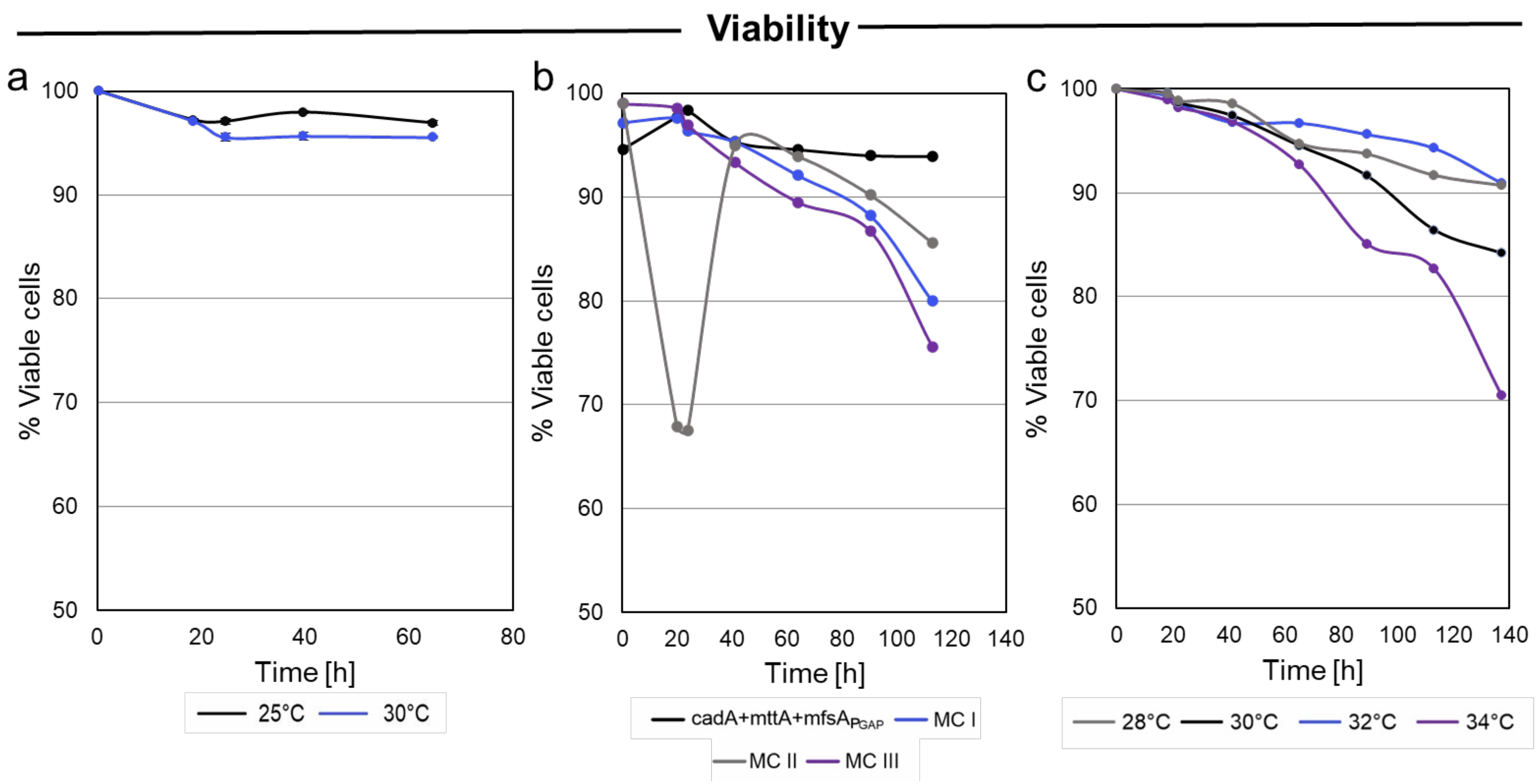


**Figure S2. Viability of strains in fed-batch cultivations. a)** The viability of the cells remained high during the fed-batch cultivation where an increased cultivation temperature of 30 °C was compared to 25 °C. In **b)** the viability of the multicopy strains (MC I, MC II and MC III) during the fed-batch cultivation at 30 °C is shown. Whilst the cadA+mttA+mfsA$_{pGAP}$ and the MC I, MC II have a stable decrease in viability over time, the MC II shows a decreased viability (> 67.5 %) at the end of the batch phase and upon methanol induction, the viability did however recover during the methanol feed phase. In **c)** the viability of the MC I strain when cultivated at 28, 30, 32 and 34 °C is shown, at 34 °C the greatest drop in viability (70.5 %) over time is experienced.

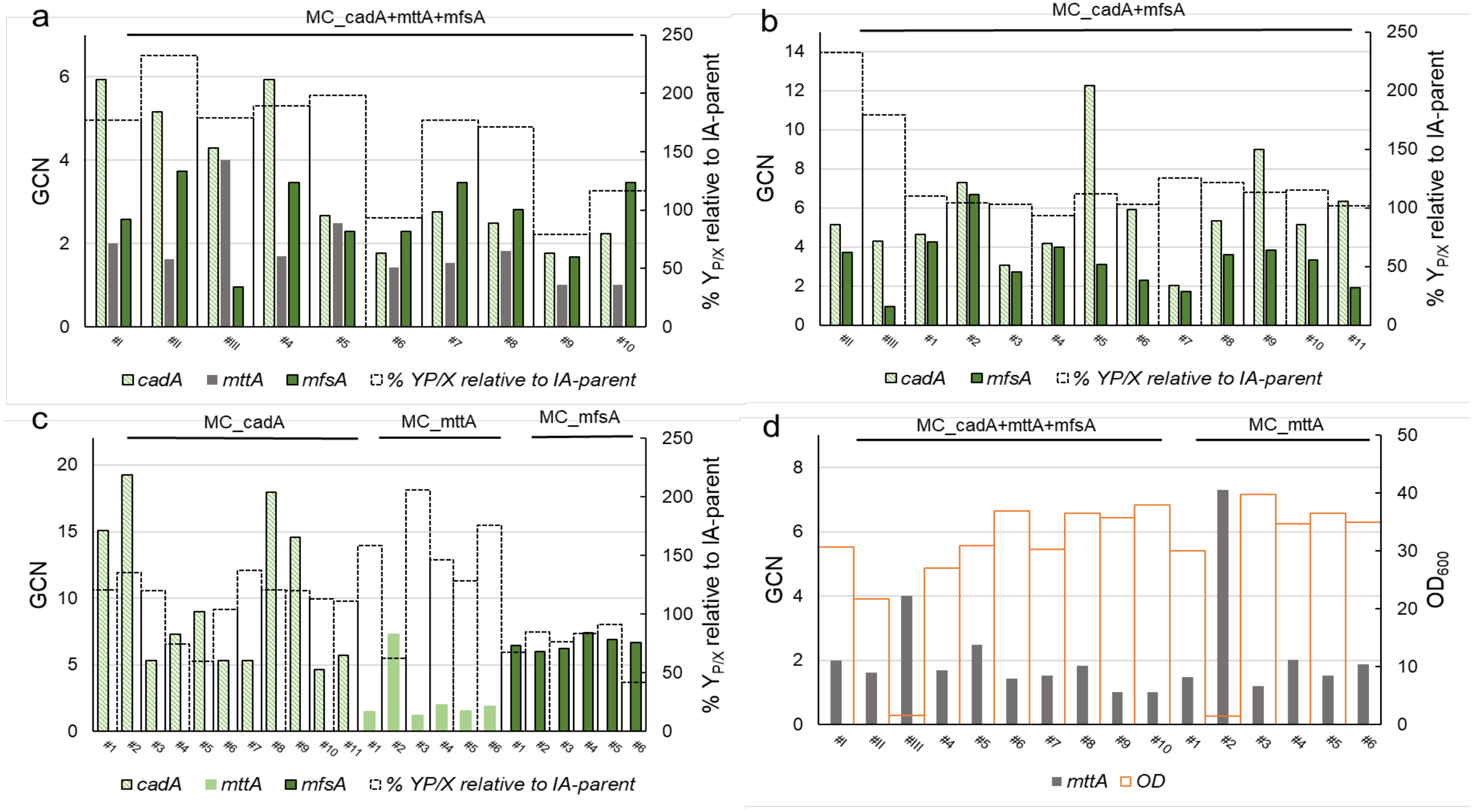


**Figure S3. The gene copy numbers (GCN) of the three heterologous genes (*cadA*, *mttA*, *mfsA*) of the generated multicopy strains (MC) was investigated via RT-qPCR relative to the cadA+mttA+mfsA$_{pGAP}$ strain**. In **a-c)** GCN of multicopy strains is shown with the relative yield obtained in the 24-deep-well plate screening in comparison to the cadA+mttA+mfsA$_{pGAP}$ strain. In **d)** the GCN of *mttA* in the MC_mttA and MC_cadA+mttA+mfsA is shown with final OD of the 24-deep well plate screening.

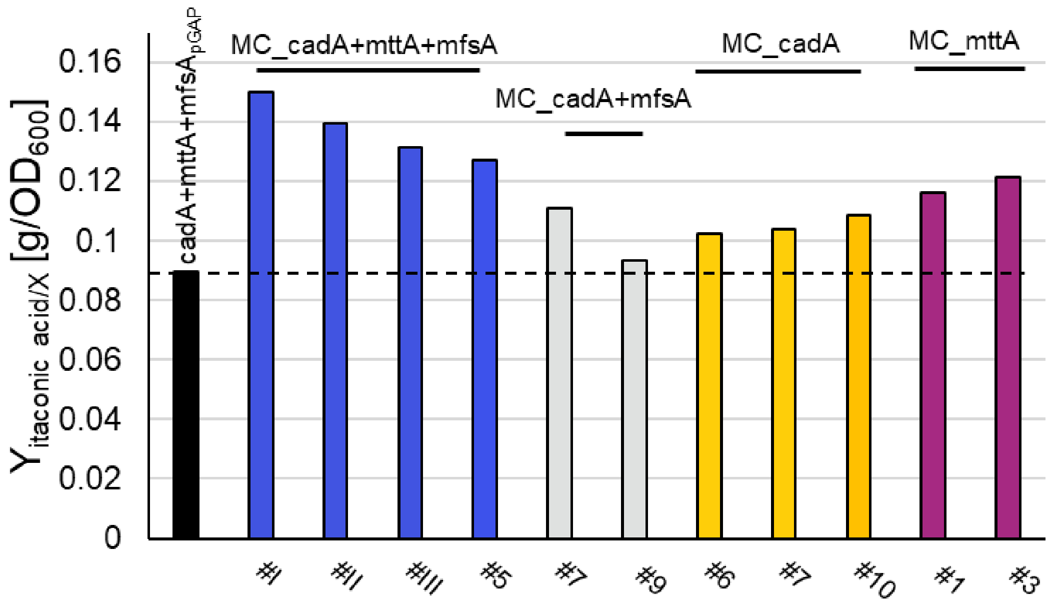


**Figure S4. Shake flask screening with multicopy strains.** A set of multicopy clones with different genotypes were selected for further investigation in a shake flask screening with the parent strain, cadA+mttA+mfsA$_{pGAP}$, as reference. The itaconic acid yields per biomass after 70 hours are shown.
